# The coculture of *Fomitopsis betulina* with *Escherichia coli* induces a nutritional stress and triggers secondary metabolism pathways

**DOI:** 10.64898/2026.09.14.751415

**Authors:** Quentin Albert, Elodie Drula, Julien Lambert, Isabelle Gimbert, David Navarro, Pierre Vilella, Marc Maresca, Attilio Di Maio, Maxime Robin, Michaël Lafond, Stephane Greff, Marie-Noëlle Rosso

## Abstract

Basidiomycete fungi are an underexplored source of specialized metabolites with significant biotechnological potential. However, the environmental cues that activate their biosynthetic pathways remain poorly understood. Here, we investigated the response of the wood-decaying fungus *Fomitopsis betulina*, to nutritional competition using co-cultures with *Escherichia coli*.

On solid medium, *F. betulina* inhibited the bacterial growth. Using a mass spectrometry-based metabolomics approach, we identified a Sumiki’s acid derivative, calcium diformate, and sulfuric acid among the compounds enriched within the inhibition zone and hypothesized that these compounds were associated with the acidification of the medium. Although *F. betulina* has previously been reported to produce the antibacterial compound piptamine, neither piptamine nor related derivatives were detected under our experimental conditions.

In liquid medium, the co-culture with *E. coli* caused the rapid depletion of the available glucose, resulting in the establishment of carbon-starvation conditions and a 44% reduction in fungal biomass. Transcriptomic analyses revealed extensive metabolic reprogramming in response to bacterial competition, including the induction of genes involved in carbon acquisition, nutrient transport, redox homeostasis, and stress adaptation. Notably, a homolog of the Velvet regulatory complex, a central regulator of fungal development and specialized metabolism, was upregulated. The co-culture induced the expression of genes associated with multiple biosynthetic gene clusters, including terpene, polyketide, and fungal RiPP.

Taken together, our results demonstrate that bacterial competition acts as a potent trigger of nutritional stress and secondary metabolism in *F. betulina*. More broadly, fungal–bacterial co-culture represents a promising alternative to extractions to identify high-value added metabolites pathways from basidiomycetes.

**Importance:** It is now well known that basidiomycete fungi are a fantastic reservoir of yet untapped secondary metabolites and antimicrobial compounds. However, the discovery of these compounds is made difficult because the genes normally remain silent, and are only expressed under specific conditions. The co-culture of fungi and bacteria is an emerging tool to decipher interactions between microorganisms and a promising alternative to chemical extractions to identify specialized metabolites from basidiomycetes. In this study, we used a bacterial competitor to induce a nutritional stress on the birch polypore, Fomitopsis betulina. We observed that the fungus responded to the stress by regulating the carbon uptake and metabolism. Carbon starvation in presence of the bacteria also triggered the over expression of a central regulator of fungal specialized metabolism, and induced the activation of terpene, polyketide, and fungal RiPP biosynthesis.

## Introduction

Saprotrophic basidiomycete fungi are of primary interest to several aspects. First, despite low growth rate, they play important roles in the carbon cycle of terrestrial ecosystems, through the decomposition of wood and other lignocellulosic materials, litter or soil organic matter, thereby contributing significantly to the global carbon cycling. Beyond their ecological significance, these fungi have attracted increasing attention as a rich source of bioactive natural products. Advances in fungal genomics have revealed a large set of biosynthetic gene clusters (BGC) with the potential to produce structurally diverse and biologically active metabolites. The potential of saprotrophic basidiomycetes as reservoirs of yet uncharacterized metabolites and their low cultivation cost make them good candidates to look for novel compounds with pharmaceutical, agricultural, or industrial applications, including antimicrobial compounds (1–6).

However, many of these biosynthesis pathways remain transcriptionally silent under laboratory conditions. Interestingly, microbial interactions are recognized as important drivers of the synthesis of secondary metabolism. Understanding the interplay between saprotrophic basidiomycetes and microbial competitors will help understand the dynamics of microbial communities in natural environments, and stimulate the production of specialized metabolites of interest. To date, most of the studies have focused on microbial interactions involving Ascomycota, particularly in the field of agricultural management, biocontrol, health and diseases (7).

For example, Ola et al. reported a 78-fold increase in the accumulation of constitutive metabolites, as well as the induction of four additional metabolites, including three undescribed natural products, in *Fusarium tricinctum* cocultured with *Bacillus subtilis* (8). Similarly, Akone et al. observed an 8-fold increase in the accumulation of a polyketide secreted by *Chaetomium sp.* during coculture with *Bacillus subtilis*, together with the induction of new metabolites, some of which exhibiting antibacterial and cytotoxic activities (9).

The identification of value-added metabolites from Fungi depends on the development of rapid mycelium-based methodologies coupled with omics approaches. For a decade, emerging methods have been developed to assess the potential of Fungi. Among them, the OSMAC approach (One Strain Many Compounds) uses mycelium growth in diverse conditions, including stress-inductive conditions to map the secondary metabolites of a strain (10, 11). Culturomics approaches have been developed with a similar strategy. Nowadays, coculture methods, in which two different organisms are grown together in the same medium, or physically separated in devices allowing chemical communication, are gaining interest, for the identification of new value-added metabolites (7, 9, 12–16), for the study of microbial cooperation during biomass degradation (17), or for ecosystem science (18–20).

However, in many coculture systems, it is difficult to decipher whether the response of the fungus is determined by nutrient depletion or by direct sensing of the microbial competitor. Fungi and bacteria exchange a wide range of signaling molecules, including quorum-sensing compounds, organic acids, volatile organic compounds, sugars, and polyols (21). As an example, Deveau et al. (18) reviewed several mechanisms driving fungi-bacteria interactions, including the response of bacteria to fungal quorum sensing molecules like farnesol, and the response of fungi to bacterial quorum sensing molecules like homoserine lactone and quinolone. Sugars, polyols, low molecular weight organic acids, and volatile organic compounds are also mentioned as chemical signals in fungi-bacteria interactions. Li et. al. further showed that the concentration of such chemicals can determine the stimulating or inhibiting effect of the signal (22).

The saprotrophs Polyporales fungi are mainly wood-specialized decomposers and provide an especially relevant model to investigate these processes. During lignocellulose degradation they secrete carbohydrate-active enzymes (CAZYmes) in the near environment of the growing hyphae, which hydrolyze the complex cellulose, hemicellulose and pectin polysaccharides into oligosaccharides or free monosaccharides such as glucose. Subsequently, the fungi may absorb the released saccharides, or conversely, the simple sugars may be retrieved by neighboring microorganisms, creating a competitive interaction for the carbon sources (17, 23, 24). In soils, some bacteria may also use fungal exudates as nutrients (25). In turn, fungi may deploy diverse defense strategies to control the bacterial growth, including the acidification of the proximal environment, the production of reactive oxygen species, the sequestration of nutrients or the secretion of antibacterial metabolites (23, 26).

In this study, we investigated the interaction between the brown-rot fungus *Fomitopsis betulina,* which was reported to produce piptamine (3, 27, 28), an antibacterial compound, and the fast growing bacterium *E. coli*. *F. betulina,* commonly referred to as the “Ice-man Fungus”, was found among the belongings of the prehistoric individual “Otzi”, and probably 5,000 years ago used for its antibacterial and antiparasitic properties (29). We hypothesized that the rapid bacterial consumption of available carbon sources would lead to nutritional stress on the fungus and trigger adaptative responses linked to carbon depletion. To test this hypothesis, we combined coculture experiments with metabolomic and transcriptomic analyses to characterize both the metabolites and the metabolic pathways underlying the fungal adaptation to bacterial competition. Our results provide new insights into the regulation of secondary metabolism in basidiomycetes and highlight coculture as a valuable strategy to explore their biosynthetic potential.

## Material and methods

### Fungal and bacterial strains

The *E. coli* strain ATCC25922 was purchased from the American Type Culture Collection (ATCC, https://www.atcc.org/, Manassas, VA, USA) and conserved at –80 °C in LB (Difco®, Sparks, Maryland, USA) with 5% (v/v) glycerol. The bacteria were subcultured in Mueller Hinton II (MHII, Difco®, Sparks, Maryland, USA) broth to reactivate the strains before use. The bacterial strains of *Enterococcus faecalis* and *Staphylococcus aureus* were preserved in our lab collection in the same conditions.

The fungal strain *F. betulina* CIRM-BRFM 860 was purchased from the Bioresource Center CIRM-CF (https://www.cirm-fungi.fr/, Marseille, France) and conserved on malt agar (Duchefa Biochemie®, Haarlem, The Netherlands) at 4 °C. It was subcultured at 25 °C on malt agar for 7 days before use.

### Confrontation assay on agar medium

A fungal plug (5 x 5 mm) was harvested from a fungal subculture and inoculated on the left side of Petri dishes with Malt Extract Agar (MEA) medium (Duchefa Biochemie®, Haarlem, The Netherlands). The cultures were incubated for 4 days at 25 °C, until the mycelium reached the middle of the plates (Fig. 1A).

**Fig. 1:**
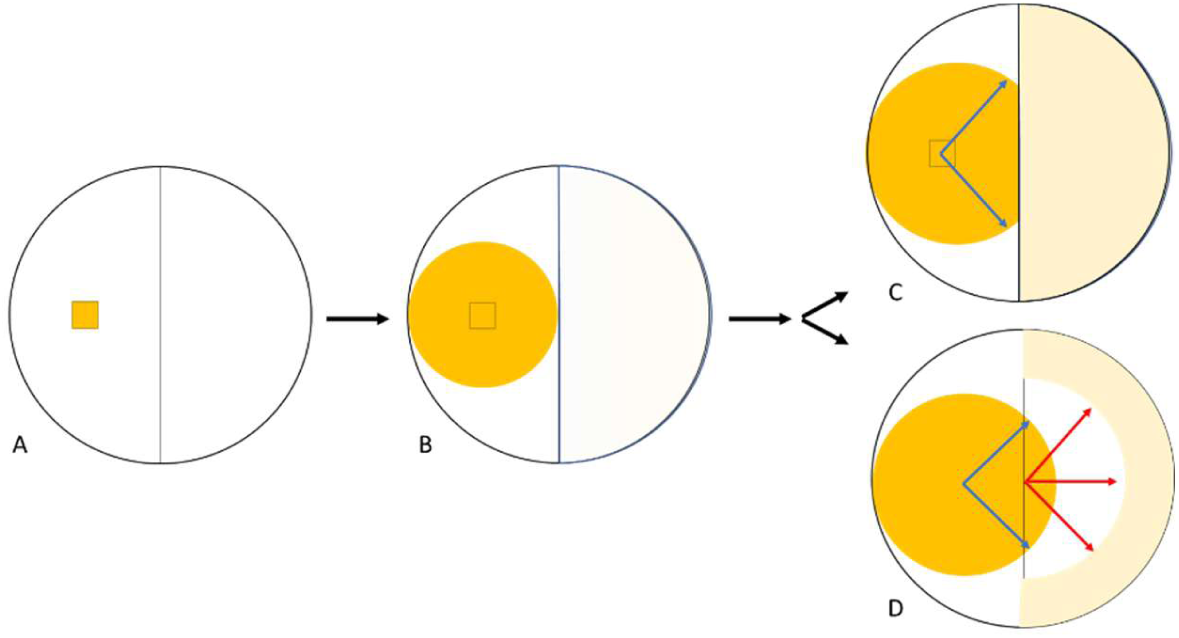
Schematic representation of the agar confrontation assay. A) Inoculation with a fungal plug (5 x 5 mm) on the left side and incubation at 25 °C. B) Inoculation with bacteria (10^c^ UFC/mL, 50 µL) and incubation 12 h at 25 °C. C) and D) Expected phenotypic observations in the absence (C) or presence (D) of antibacterial activity. The blue arrows illustrate the fungal growth, and the red arrows show the bacterial growth inhibition zone.

Then, 50 µL of a normalized bacteria inoculum (OD_600_ = 0.1, corresponding to 10^6^ UFC.mL^-1^) was spread on the surface of the right side of the plates (Fig. 1B). The plates were incubated at 25 °C for 12 h. Plates inoculated with the fungus alone or bacteria alone were used as controls.

The pH on the surface of the agar medium was measured using an InLab Pro Surface probe (Metler-Toledo®, Greifensee, Switzerland). The assay and the controls were done in 5 replicates.

### Metabolomic study and metabolite characterization in the agar confrontation assay

#### Sampling

Samples were collected with the base of Pasteur pipettes on three different zones of the agar plate (five samples per zone): beneath the fungus, beside the fungus (in the inhibition zone) and in the bacterial mat (see Supp File). Pure culture of fungi served as controls. Each zone was extracted with 5 mL MeOH (LC/MS grade, Carlo Erba®, Peypin, France) under ultrasonication during 10 min at 40 °C (J.P. Selecta, Barcelona, Spain). The supernatants were then transferred into 20 mL vials, dried and recovered in 1 mL MeOH before UHPLC-HRMS analysis.

#### UHPLC-HRMS Metabolomics & Multivariate analyses

Mass spectrometry metabolomic analysis was performed on an Ultra High-Performance Liquid Chromatography (UHPLC Ultimate 3000RS, Thermo®, Courtaboeuf, France) coupled to a High-Resolution mass spectrometer (ESI-qToF Impact II, Bruker®, Wissembourg, France). Samples were eluted on a Polar C_18_ column (150 x 2.1 mm, 1.6 *µ*m, Phenomenex®, Le Pecq, France) maintained at 42 °C using a solvent gradient composed of water (solvent A) and acetonitrile (solvent B), both supplemented with 0.1% formic acid (all solvents were LC/MS grade, Carlo Erba®, details in Raw Data, see Method and Data Supp file, sheet 2_LCMS Metabolomics). The data were acquired in positive mode and processed using MZmine (version 4.6.1) for chromatographic and spectral deconvolution, leading to the production of a data matrix containing 3,255 chemical features. After manual curation, the data matrix (2,337 features) was uploaded onto MetaboAnalyst (Version 6.0) for multivariate data analyses. After filtration on the platform, the data matrix (463 features) enabled a selection of significant chemical features corresponding to potential fungal metabolites (biomarkers). MS/MS spectral annotations for the putative identification of these biomarkers were performed on Bruker Compass DataAnalysis by comparing the data with those available in databases (e.g. Mass Bank of Europe) or by complementing the annotations using *in silico* MS/MS annotation tools (SIRIUS version 6.2.2 and MetFrag).

### Fungal metabolite production, purification and characterization

A process for metabolite production and purification was set up to structurally identify a metabolite detected as biomarker (biomarker#6) detected in the agar confrontation.

#### Metabolite production

The fungus was first pre-cultivated on static, liquid malt extract broth (Duchefa Biochemie®, Haarlem, The Netherlands) until the development of the fungal mat (25 °C, 15 days). The fungal mat was then filtered and fragmented with a T25 digital Ultra Turrax® disperser (Ika®, Staufen, Germany). The fragmented mycelium was used to inoculate 1 L of malt extract broth (MEB, Duchefa Biochemie®, Haarlem, The Netherlands) in an Erlenmeyer flask maintained under stirring (25 °C, 120 rpm) for three weeks. The supernatant was filtered with a 10 kDa cutoff membrane (Vivaflow 200 Flip-Flop, Sartorius®, Göttingen, Germany) with a peristaltic pump (503U Watson-Marlow®, Cheltenham, England, UK), to retrieve the metabolites. A final freeze-drying step led to the obtention of 8.17 g of dry secretome.

#### Metabolite purification and characterization

The dry secretome was solubilized in H_2_O (UHPLC/MS grade, Carlo Erba®) at a concentration of 250 mg/mL. The metabolites were purified by semi-preparative chromatography on a PLC2050 preparative LC system (Gilson® Purification S.A.S., Saint-Ave, France) equipped with a PDA detector. The solution was injected (1 mL/ injection) on a Polar C_18_ column (Luna Omega, 250 x 10 mm, 5 *μ*m, Phenomenex®) and eluted at room temperature with H_2_O (MilliQ®, Select HP80, Purite Ltd, Thame, United-Kingdom) and acetonitrile (preparative HPLC grade, Carlo Erba®), both supplemented with 0.1% (v/v) formic acid (LC/MS grade, Carlo Erba®) at a flow rate of 4 mL/min (see gradient details in details in Raw Data Method and Data Supp file). The compounds were detected at 270 nm, and the fractions were automatically collected every 1.5 min (6 mL/tube). The collected fractions were dried under vacuo (MiVac Quattro from Genevac®, Biopharma technologies France, Diémoz, France). Metabolite 1 (retention time rt = 10.05 min) and metabolite 2 (rt = 10.42 min) were purified and further characterized by HRMS/MS in both positive and negative modes as well as ^1^H NMR (Bruker Avance II, 600 MHz, 1.6 and 2.0 mg for metabolite 1 and 2 resp. in 300 µL D_2_O, analysis of 80 µL in 2 mm tubes).

### Synthesis of piptamine and piptamine derivatives

To synthesized N-Methyl-N-pentadecylbenzenemethanamine (Piptamine), a mixture of N-Benzylmethylamine (10 mmoles), 1-Bromopentadecane (10 mmoles) and K_2_CO_3_ (20 mmoles) were solubilized in CH_3_CN (20 mL) and heat at 50 °C for 18 h. The resulting mixture was filtered off and washed with CH_3_CN (20 mL). CH_3_CN was removed under vacuum and the resulting oil was solubilized in dichloromethane (20 mL) and then washed with brine (20 mL). The organic phase was then purified by flash chromatography (DCM/MeOH 9/1) to give the pure N-Methyl-N-pentadecylbenzenemethanamine as a light-yellow oil. Yield 90%, ^1^H NMR (300 MHz, CHLOROFORM-d) δ ppm: 0.82 – 0.89 (t, J= 6.42 Hz, 3 H, CH_3_), 1.24 (s, 24 H, CH_2_), 1.49 (q, 2 H, CH_2_), 2.15 (s, 3 H, N-CH_3_), 2.28 –2.37 (t, J= 7.61 Hz, 2 H, N-CH_2_-CH_2_), 3.44 (s, 2 H, CH_2_benzyl), 7.14 – 7.32 (m, 5 H, C-Hbenzyl). ^13^CNMR (75 MHz, CHLOROFORM-d) δ ppm: 14.08 (CH_3-15_), 22.66 (CH_2-14_), 27.39 (CH_2-3_), 27.44 (CH_2-2_), 29.35 (CH_2-4_), 29.58 (CH_2-12_), 29.63 (CH_2_),29.68 (CH_2_), 31.91 (CH_2-13_), 42.18 (N-CH_3_), 57.59 (CH_2-1_), 62.31 (N-CH_2bn_), 126.76 (C-4), 128.08 (C-3,C-5), 128.99 (C-2,C-6), 139.24 (C-1). Anal. Calcd. for C_23_H_41_N: C, 83.31; H, 12.46; N, 4.22. Found: C, 83.22; H, 12.51; N, 4.09. *m/z*: 331.3239 (100.0%), 332.3273 (24.9%), 333.3306 (3.0%). All the piptamine derivatives have been synthetized by the same way by varying the halogenoside chain.

### Antibacterial activity assays

The antibacterial activity was quantified using the agar diffusion method and the serial dilution method. We determined the minimal inhibition concentration (MIC) against *E. coli* ATCC 25922, *Enterococcus faecalis* or *Staphylococcus aureus* following the EUCCAST recommendations. Briefly, *E. coli* ATCC 25922, *E. faecalis*, and *S. aureus*, were grown overnight in MHII broth (30 °C), and 100 µL of a standardized inoculum (OD_600_ = 0.1) were spread on fresh MHII agar medium. Then, the positive control (Ampicillin disk 5 µg, Oxoid®), 5 µL of pure synthetized piptamine, or 5 µL of secretome (100 mg/mL) or 5 µL of purified metabolite 1 (1 mg/mL) were spotted on the bacterial mat. The Petri dishes were incubated at 30 °C overnight before measurement of the inhibition halo. The serial dilution method was performed in 96-well plates (Greiner Bio-One®, Kremsmünster, Oberösterreich) using dilution series of secretome ranging from 0.1 to 25 mg/mL in sterile water, against a standardized *E. coli* inoculum (OD_600_ = 0.1 corresponding to 10^5-6^ UFC/mL). Each assay was performed in four replicates. The results obtained with the secretome are mentioned as “MIC-like” in the following sections.

### Coculture experiment in liquid medium

Three fungal plug (5 x 5 mm) were harvested from a fungal subculture on MEA and used to inoculate 100 mL of MEB. The fungus was first cultured in MEB at 30 °C, 120 rpm alone. After 3 days, we added 1 mL of a standardized inoculum of bacteria (OD_600_ = 0.1) precultured on MHII broth. The cocultures were further incubated at 30 °C, 120 rpm. Pure cultures of the fungus and pure cultures of bacteria in the same conditions were used as controls. After 3 days, the mycelium was recovered on a sterile pad, rinsed with 25 mL NaCl 0.09%. The mycelium was dried at 110 °C to quantify the fungal biomass (n = 5), freeze-dried until extraction of total RNAs (n = 3) or transferred on fresh MEA plates to test fungal survival (n = 5, incubation at 25 °C). To recover the bacteria, the spent culture medium was centrifuged at 7,000 rpm for 5 min. The bacteria were resuspended in 5 mL NaCl 0.09% and spread on fresh MHII agar (10 µL) to perform the survival test (30 °C overnight).

The glucose concentration and pH (Accumet 13-620-108-B, Fisher Scientific®, Pittsburgh, Pennsylvania, USA) were monitored during the co-culture experiment. Briefly, 1 mL of spent culture medium was sampled each day. We used the Glucosepod-400 enzymatic assay (Libios®, Vindry Sur Turdine, France) to quantify glucose, following the recommendations of the manufacturer.

### RNA extraction, sequencing, and analyses

For each condition, RNA was extracted using 100 mg of ground mycelium and 1 mL TRIZOL (Ambion®, Carlsbad, California, USA). RNAs were precipitated with isopropanol (Sigma-Aldrich®, Burlington, Massachusetts, USA), treated with DNAse I (QIAGEN®, Hilden, Germany) and resuspended in 25 μL RNAse Free water. RNA purity and integrity were analyzed on NanoDrop® Spectrophotometer and Agilent Technologies® 2100 BioAnalyzer (Santa Clara, California, USA). The RNA libraries were built using Illumina® stranded mRNA Prep and sequenced on NovaSeq® 6000 (2 x 150 bp) at the GeT-PlaGe France Genomique platform. The sequences were mapped on the genome of *F. betulina* CIRM-BRFM 1772 (https://mycocosm.jgi.doe.gov/Pipbet1_1/Pipbet1_1.home.html; Genbank accession GCA_022606075.1). The RNASeq analysis was performed at the GeT-Biopuces platform using the R software (30), Bioconductor (31) packages including DESeq2 (32, 33), and the SARTools package developed at PF2 – Institut Pasteur (34). The genes with normalized basemean read counts < 5 were not considered in the analysis. Genes were considered differentially regulated if |log2fold change| ≥ 2 and p adj < 0.05.

The protein sequences were retrieved from Mycocosm (35) and the predicted functions were assigned by 1) searching for Pfam domains, using the InterPro database (https://www.ebi.ac.uk/interpro) (36, 37), 2) expert annotation of CAZymes (38), 3) annotation of secondary metabolite gene clusters using antiSMASH (39), and 4) annotation of transmembrane domains using TMHMM annotations from InterPro. Signal P 5.0 was used to predict the presence of putative secretion signal peptides with default parameters (https://services.healthtech.dtu.dk/services/SignalP-5.0/). For some proteins, the annotations were completed by BLASTp against SwissProt (https://blast.ncbi.nlm.nih.gov/Blast.cgi).

The KEGG identifiers were searched for *F. betulina* predicted protein sequences using the KEGG Automatic Annotation Server Ver. 2.1 (40) with the GHOSTZ method on a set of 40 genomes including eukaryote model organisms and fungi.

### Accessibility of the data

The raw data that support the findings of this study are available from the corresponding author upon reasonable request.

The raw data from metabolomic study are available on zenodo (10.5281/zenodo.20283660). (Metabolomics data can be viewed by the reviewers with this link: https://nmrxiv.org/project/F8RBTH8GFjrruQCC68NOXHxn0K2TttgoMrBiHrYC). Data of Sumiki’s acid calcium complex (Metabolite 1) were deposited on NMRxiv for NMR acquisitions (https://doi.org/10.57992/nmrxiv.p175) and MassiVe for MS acquisitions (MS^2^ spectra acquired in positive mode at different collision energies: from <u>CCMSLIB00017469757</u> to <u>CCMSLIB00017469762</u>; MS^2^ spectrum acquired in negative mode: <u>CCMSLIB00017469763</u>).

The raw data from transcriptomic study are available in the ArrayExpress database (http://www.ebi.ac.uk/arrayexpress) under accession number E-MTAB-17615.

Tanscriptomics data can be viewed by the reviewers with this link: https://www.ebi.ac.uk/biostudies/ArrayExpress/studies/E-MTAB-17615?key=672724dc-7fe3-4762-8fa5-812e68397c96

## Results and discussion

### Fomitopsis betulina exhibits antibacterial activity on agar confrontation assays

We first confirmed that the strain *F. betulina* CIRM-BRFM 860 had antibacterial activity using cocultures of the fungus and *E. coli* ATCC 25922 on agar plates. We observed a clear inhibition of the bacterial growth at distance from the mycelium (Fig. 2).

**Fig. 2:**
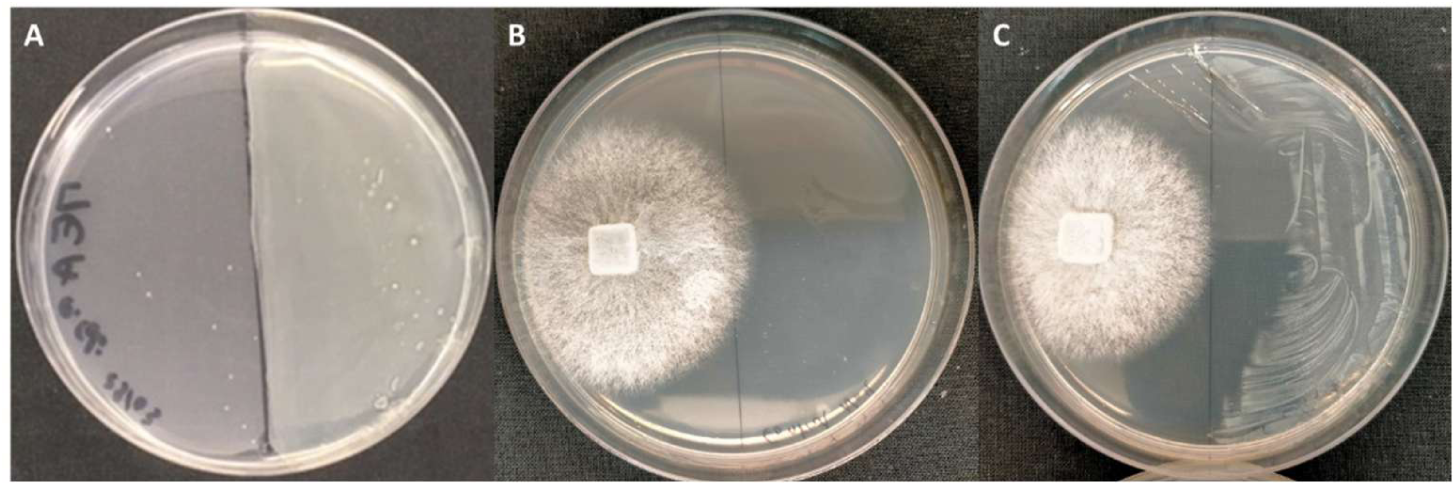
Coculture on MEA medium. A) Pure culture of E. coli ATCC 25922 (overnight, 25°C), B) Pure culture of F. betulina CIRM-BRFM 8c0 (4 days, 25°C), C) Coculture of F. betulina CIRM-BRFM 860 and E. coli ATCC25922 (4 days, 25°C).

In a previous study, Schlegel and collaborators identified piptamine in the spent culture medium of *F. betulina* Lu-9-1, and showed its antibacterial activity against several Gram positive bacteria, including *Staphylococcus aureus* SG 511 (0.78 µg/mL), *Bacillus subtilis* ATCC6633 (1 µg/mL), and *Enterococcus faecalis* 1528 (1.56 µg/mL), as against Gram negative bacteria like *Escherichia coli* SG 458 (12.5 µg/mL) (28). In order to test if piptamine was produced by *F. betulina* CIRM-BRFM 860 in our conditions, we synthetized the molecule and used it as a standard in UHPLC experiments. *F. betulina* CIRM-BRFM 860 did not produced piptamine in our experimental conditions (neither in pure culture nor after coculture). Surprisingly, the synthetized piptamine failed to show any antibacterial activity against *E. coli* ATCC 25922 *S. aureus*, and *E*. *faecalis.* Because the piptamine could be chemically modified by the fungus in our conditions, we also synthetized piptamine derivatives and used them as standards in the UHPLC experiments. Yet, we did not identify the piptamine derivatives among the secreted metabolites (Fig. 1 Supp Data).

In a search for the molecule(s) responsible for the observed antibacterial activity of *F. betulina* CIRM-BRFM 860, we developed a MS-based metabolomic analysis from the agar confrontation assay. We sampled the agar medium beneath the fungal mycelium, nearby the growing mycelium, in the inhibition zone between the fungus and the bacterial colony and beneath the bacterial colony. Pure cultures of mycelium and sterile agar medium were used as controls. The data matrix obtained in positive mode contained a total of 2,337 features. After filtration on MetaboAnalyst (filtering features based on technical repeatability of QC samples (20%), and Interquantile range filtering (40%), see details in Raw Data, Method and Data Supp file sheets 6 to 9), a total of 463 features remained. The data were then Log-transformed and normalized (range scale). The Principal Component Analysis highlighted a first differentiation along PC1 axis accounting for 43.8% of the variance (Fig. 4A). Control samples (agar medium, mycelium) grouped together. A second group along this axis was formed by media extracts collected beneath the fungus (alone or in coculture). The extracts from the inhibition zone were clearly separated from the other samples along PC2 (24.8% of the variance).

**Fig. 3:**
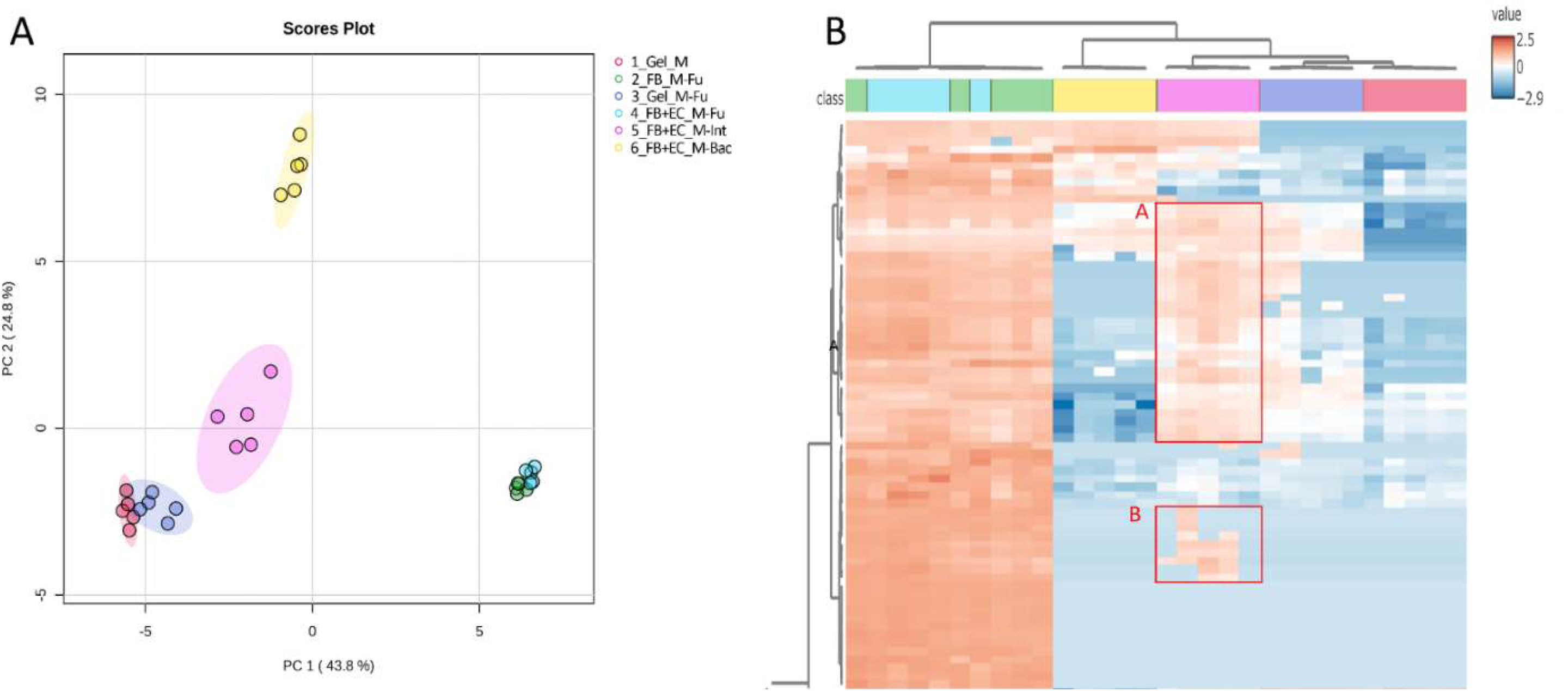
A) Principal Component Analysis of metabolomic data acquired in positive mode by Ultra High-Performance Liquid Chromatography coupled to a High-Resolution Mass Spectrometer, and B) heatmap highlighting overrepresented features in the inhibition zone. The features detected in the inhibition zone (groups A and B) were also found beneath the fungus (alone or in co-culture), not in the control agar media and less-represented beneath the bacterial colony. 1_Gel_M: agar media (control), 2_FB_M_Fu: agar media under the fungus (alone), 3_Gel_M_Fu: agar media nearby the fungus (alone), 4_FB+EC_Fu: agar media under the fungus (coculture), 5_FB+EC_Int: agar media in the inhibition zone between the fungus and the bacterial colony, 6_FB+EC_M-Bac: agar media under the bacterial colony (coculture).

**Fig. 4:**
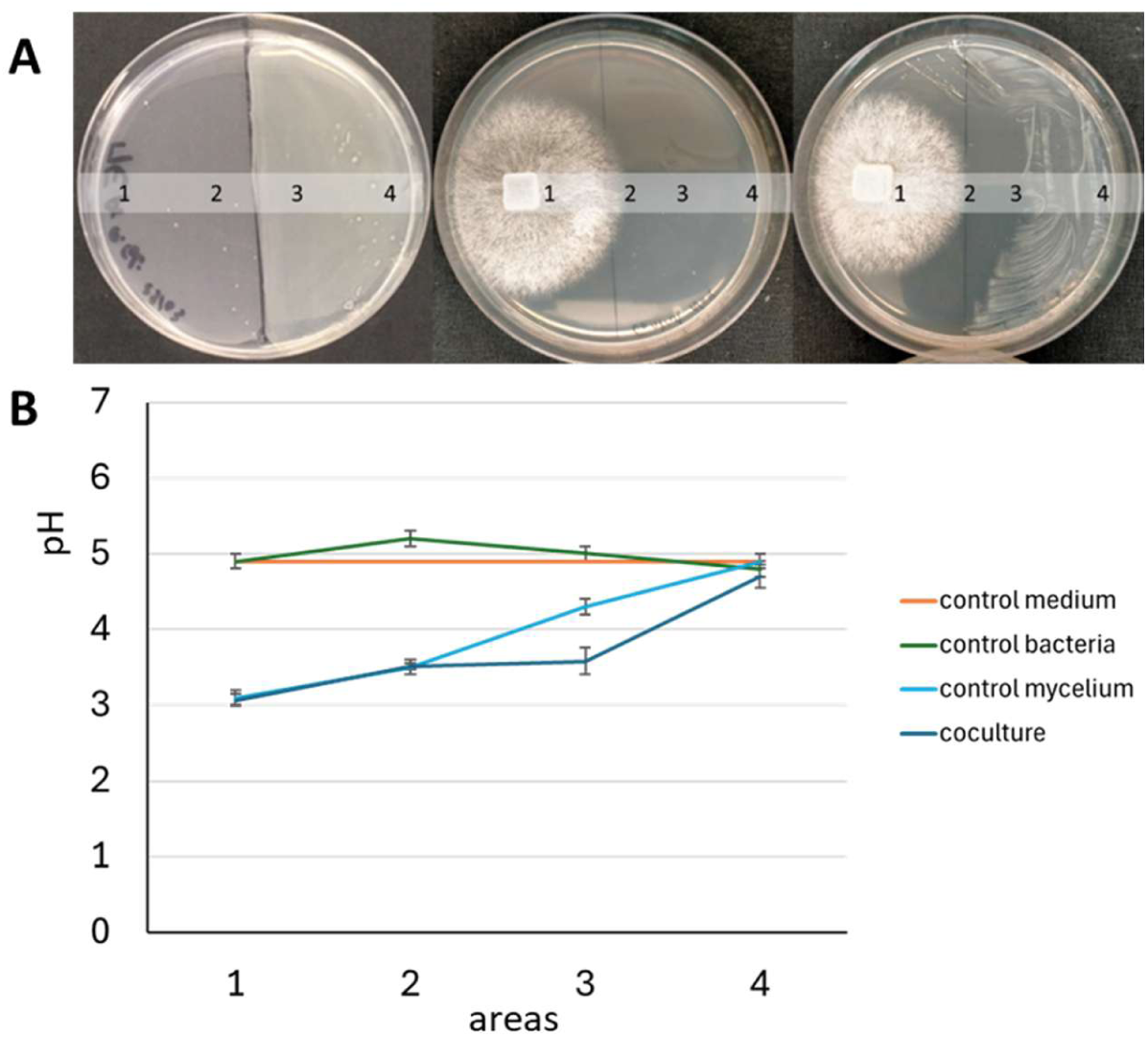
pH measurement on the agar assay of Fomitopsis betulina CIRM-BRFM 8c0 coculture with E. coli ATCC 25S22. *A) from left to right: photograph of agar medium inoculated with the bacteria only, the fungus only, and both the bacteria and the fungus. The pH was measured on the surface of the agar in areas 1 to 4. B) pH measurements on the surface of the agar in areas 1 to 4 (n = 4 in each condition)*.

Based on the heatmap analysis, we identified 38 features that were overrepresented in the inhibition zone (Fig. 4B groups A and B), which were also detected in the agar medium beneath the fungus (alone or in co-culture). The features were grouped by molecular identity, and their spectra were further annotated leading to the putative identification of 18 metabolites, among which 10 were amino-acid derivatives (details in Raw Data, see Method and Data Supp file sheet 11 and 12_ LCMS_Biomarker table concat). One selected metabolite, designated biomarker #6 (= isolated metabolite 1), was associated with a cluster of six features corresponding to in-source fragments and exhibited a MS spectrum that was initially difficult to interpret. Nevertheless, this biomarker attracted our attention because it was clearly detectable at 270 nm, suggesting the presence of aromatic chromophore(s) facilitating also the development of a semi-preparative chromatographic method for its purification using UV monitoring.

The biomarker#6 was identified as a Sumiki’s acid calcium complex (three units of Sumiki’s acid in complex with Ca^2+^, C_18_H_16_CaO_12_), which was overrepresented in the growth inhibition zone, as compared to both the mycelium and the bacterial zone (details in Raw Data, see Method and Data Supp file sheet 11_LCMS_Biomarker study, Biomarker #6 and 13_MS_NMR_Biomarker 6 ID for structural elucidation).

The Sumiki’s acid is a furan-derived Low Molecular Weight Organic Acid (LMWOA; 5-(hydroxymethyl)furan-2-carboxylic acid, CAS: 6338-41-6) (41, 42). Fungi are well known producers of LMWOAs, which often are siderophores. Fungal LMWOAs and siderophores play a key role in the mineral nutrition and metal biosorption. We hypothesized that the Sumiki’s acid could be secreted by the fungus to scavenge minerals, such as Ca^2+^, or Fe^2+^, and deprive the bacteria from it (18, 43). The Sumiki’s acid calcium complex did not reveal any antibacterial activity (1 mg/mL in purified water, agar diffusion method) against *E. coli* ATCC25922 or *Staphylococcus aureus*.

During biomarker#6 purification, we were able to isolate another compound (called Biomarker#6 congener = metabolite 2, see 3_Preparative chromatography, 14_MS_Congener of Biomarker 6) that eluted just after Sumiki’s acid calcium complex. We were unable to structurally characterize this compound, but by injecting it on UHPLC at 100 *µ*g/mL, we recovered Sumiki’s acid Calcium complex and two highly polar compounds that could be annotated as calcium diformate and sulfuric acid. Sumiki’s acid Ca complex (metabolite 1) seems to be produced from its congener that is originally present in the agar media. One or both organisms may bio-transform the congener in Sumiki’s acid Ca complex liberating acidifying compounds in the media.

These acids are likely responsible for the pH gradient measured in Petri dishes (Fig. 5). The pH on the agar assays ranged from 3 at the mycelium inoculation plug to 5, which is the native pH of the medium, at distance from the growing mycelium. Interestingly, we observed that the pH was more acidic in the bacterial growth inhibition zone compared to the same area in the absence of bacteria, hypothesizing that the acidification might be increased, or induced, in the presence of the bacteria.

**Fig. 5:**
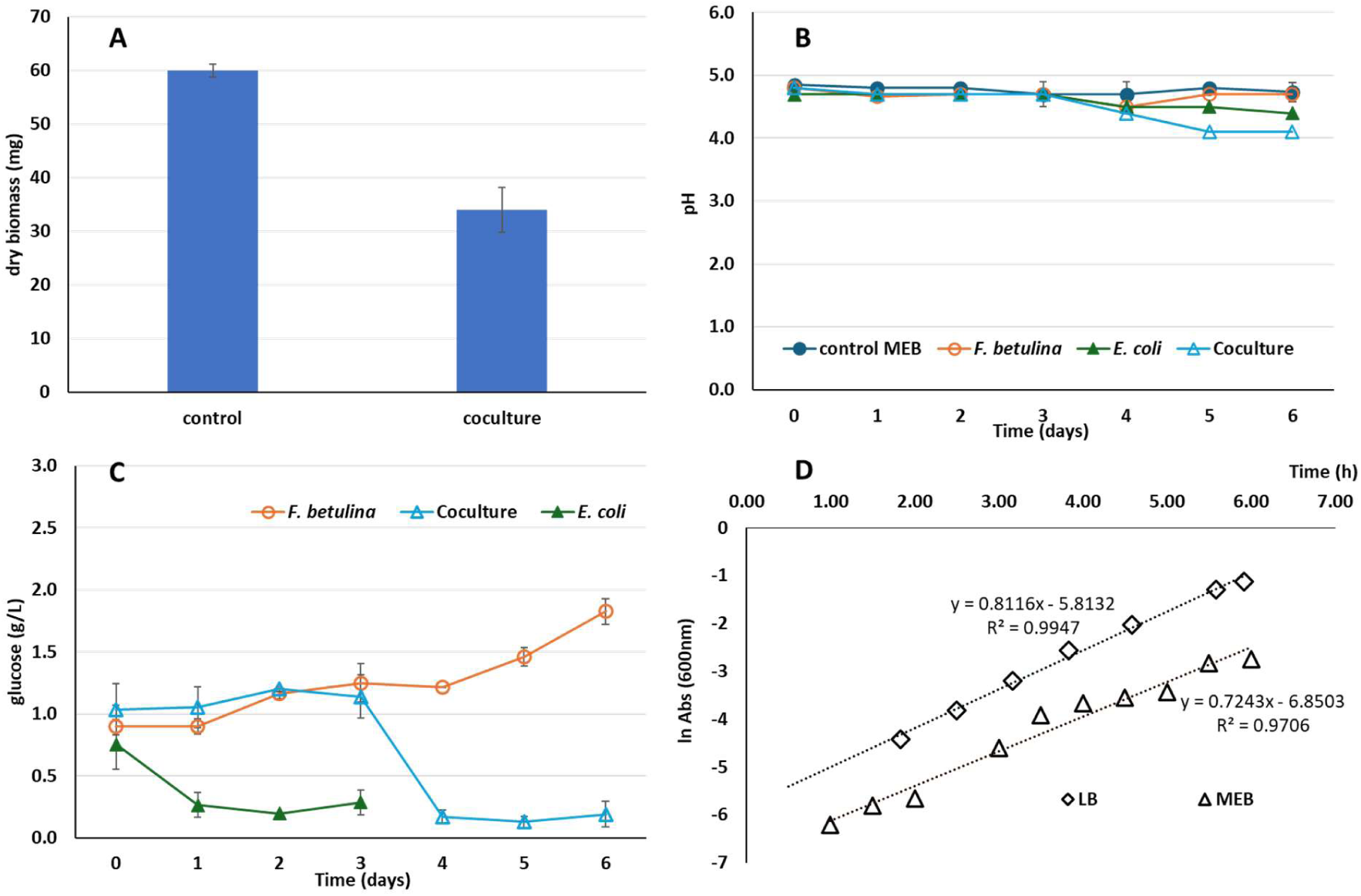
Phenotypic characterization of the coculture in liquid medium. A) Production of fungal dry biomass; B) evolution of the pH over time (days); C) evolution of the glucose concentration (g/L) over time (days), and D) growth curve of E. coli on LB and MEB.

In conclusion, the antibacterial activity we observed in *F. betulina* CIRM-BRFM 860 could be a consequence of the acidification of the medium by the growing mycelium, rather than a consequence of diffusible antibacterial metabolites, or mineral nutrition competition. This conclusion is relevant with the mentioned acidification by other studies and with the niche colonization strategy of brown-rot fungi such as *F. betulina* (18, 23).

### The fungus and bacteria competed for nutrients during the coculture

We tested whether the fungus and the bacteria would compete for the carbon source during the coculture in liquid medium. We first cultured the fungus in malt extract broth (MEB) for 3 days, after which we added 1 mL of a standardized inoculum of bacteria. The cocultures were further incubated for 3 days before analysis.

After 6 days growth in control pure cultures, *F. betulina* produced 60 mg biomass (dry weight), the pH remained stable and the glucose concentration increased over time, as a result of the expected maltose and starch degradation by the fungal maltase (Fig. 6A-C). This result suggests that the fungus degraded maltose faster than it absorbed glucose, as described in the literature (44). We verified the bacterial growth rate in pure cultures and observed a slightly reduced growth rate in MEB as compared to the growth rate in LB (Fig. 6D).

**Fig. 6:**
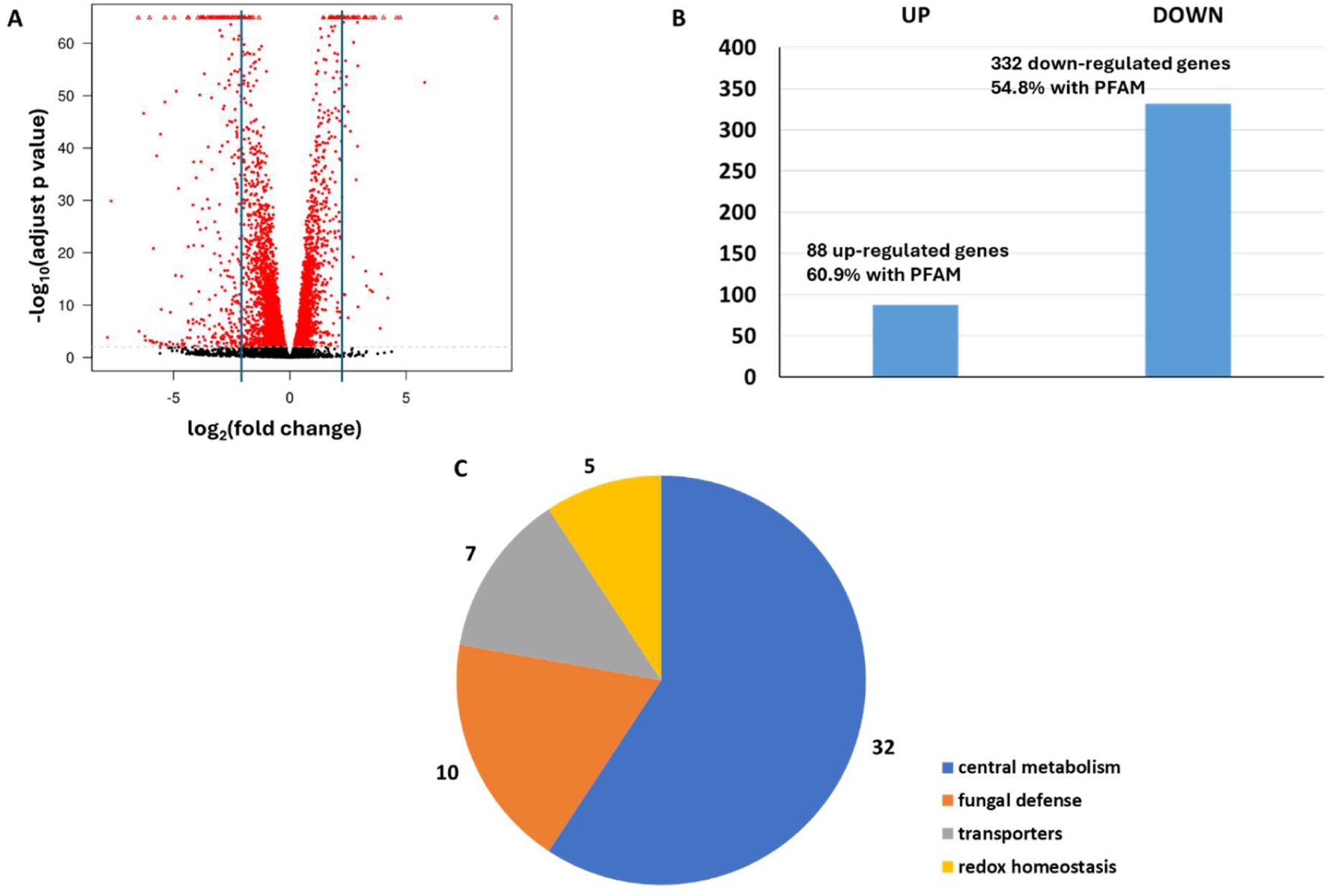
Overview of transcriptome regulation during the coculture. A) Volcano plot showing the differential transcription of the genes. The differentially expressed genes are shown in red. For this study, we focused on the differentially expressed genes with |log_2_ (fold-change) | ≥ 2, corresponding to a 4 time up-or down-regulation of the transcription. B) Counts of up-and down-regulated genes in the coculture compared to the control, C) up-regulated genes among general metabolic pathways.

After3 days of growth in coculture with *E. coli*, the growth of *F. betulina* was reduced to 34 mg dry weight (56%) (Fig. 6A). After the bacteria were added to the culture (day 3), the glucose concentration was reduced to zero within 1 day, similar to the glucose consumption observed in bacterial pure cultures (Fig. 6C).

The bacterial count on agar medium at the end of the experiment did not reveal any impact of the coculture on the bacterial growth (10^9^ UFC/mL in both the control pure culture and the coculture).

Conversely, the rapid consumption of the glucose by the bacteria and the drastic reduction of the fungal growth revealed that *E. coli* was an efficient nutrient competitor for *F. betulina* in these experimental conditions.

### Carbon depletion during the coculture induced a metabolic stress in *F. betulina*

We analyzed gene regulation in *F. betulina* after 3 days of coculture using RNA sequencing. From the 5 biological replicates of *F. betulina* grown on pure culture, we obtained a mean of 113 M reads per sample. From the 3 replicates in coculture, we obtained a mean of 156 M reads. Among them, 84% had a unique match on one of the 12,313 genes from the genome of *F. betulina* CIRM-BRFM 1772 (Genbank accession GCA_022606075.1). In total, 11,127 genes (90%) were transcribed in the tested conditions (mean normalized read counts > 5). We filtered the genes that were differentially expressed between the coculture and the fungal pure culture (|log2 fold change| ≥ 2, with adjusted *p* value < 0.05). As shown in Fig. 7A and B, we identified 88 up-regulated and 332 down-regulated genes.

**Fig. 7:**
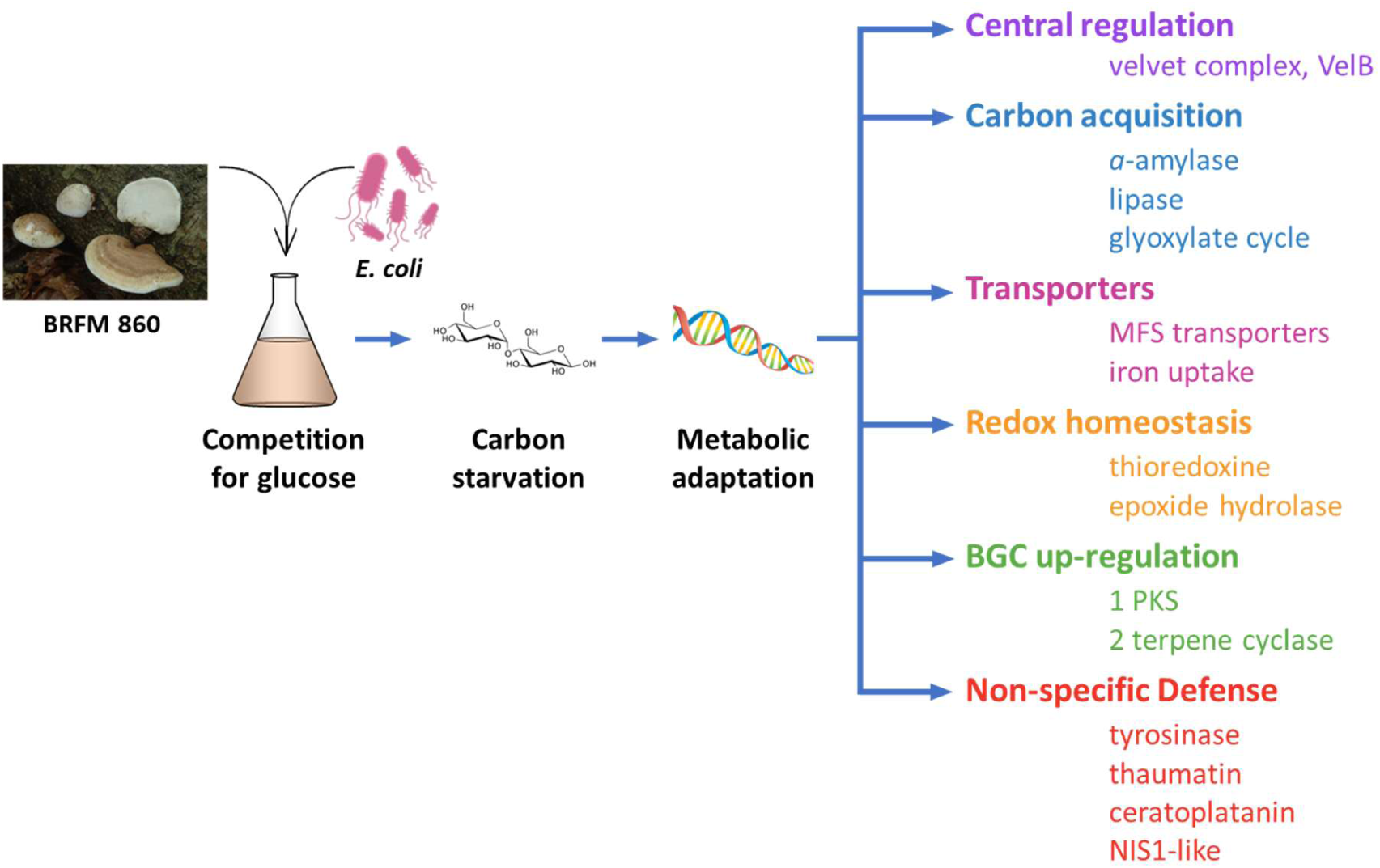
Overview of the metabolic adaptation of F. betulina BRFM 8c0 to bacterial competitor E. coli ATCC25S22.

Most of the down-regulated genes (230/332) lacked functional annotations. Considering the KOGG Class, 26 genes had “General functions prediction”, 20 were related to “Energy production and conversion”, and 18 were related to “Secondary metabolites biosynthesis, transport and catabolism”. The three most down-regulated genes with predicted function were related to lipid transport and metabolism (homolog of ProtID #862698 in *F. betulina* CIRM-BRFM 1772, repression factor –92), and energy production and conversion (homolog of ProtID #787022 and #937870, repression factor –79 and –53, respectively). We found eight down regulated genes potentially related to ROS homeostasis (homolog of ProtID #796660, #887663, # 821297, #857028, #874616, #853858, #200648, and #194970) with repression factors ranging from –18 to –5, suggesting that the presence of the bacteria disturbed the redox homeostasis of the fungus.

Among the 88 up-regulated genes, 53 (60%) had a Pfam domain and 31 (35.6%) were predicted to code for secreted proteins (Fig.7C). Regarding central metabolism, we found only one CAZYme, an alpha-amylase (ProtID #10028) from the GH13 family (www.cazy.org; (38)), that was up-regulated by a factor 5 in the coculture as compared to the pure culture (Table 1; sup data). This result showed that the fungus responded to carbon depletion in the culture medium to enhance carbon acquisition from the malt extract. GH13could also be involved in the Fungal Cell Wall remodeling (45). We found a secreted lipase 3 (ProtID #190129, 34% identity with a lipase from *Malassezia restricta* CBS 7877, a Basidiomycota yeast, accession number: A8PUY5.1). Lipases are used to hydrolyze organic compounds as alternative carbon sources when carbohydrates are not available to produce energy. In addition, we found two genes related to glutamine and nitrogen metabolism (ProtID #879792 and #769095), which could regulate nitrogen homeostasis in this stress conditions.

**Table 1:**
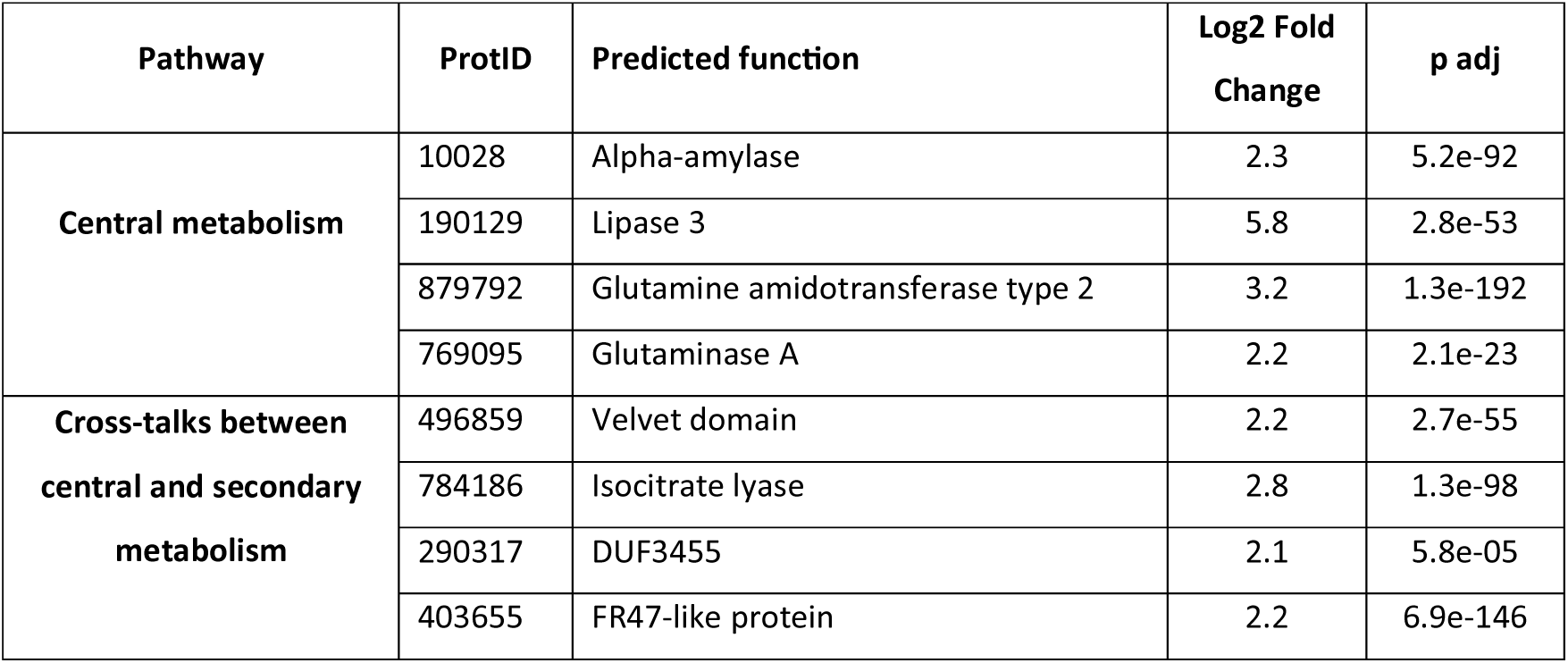
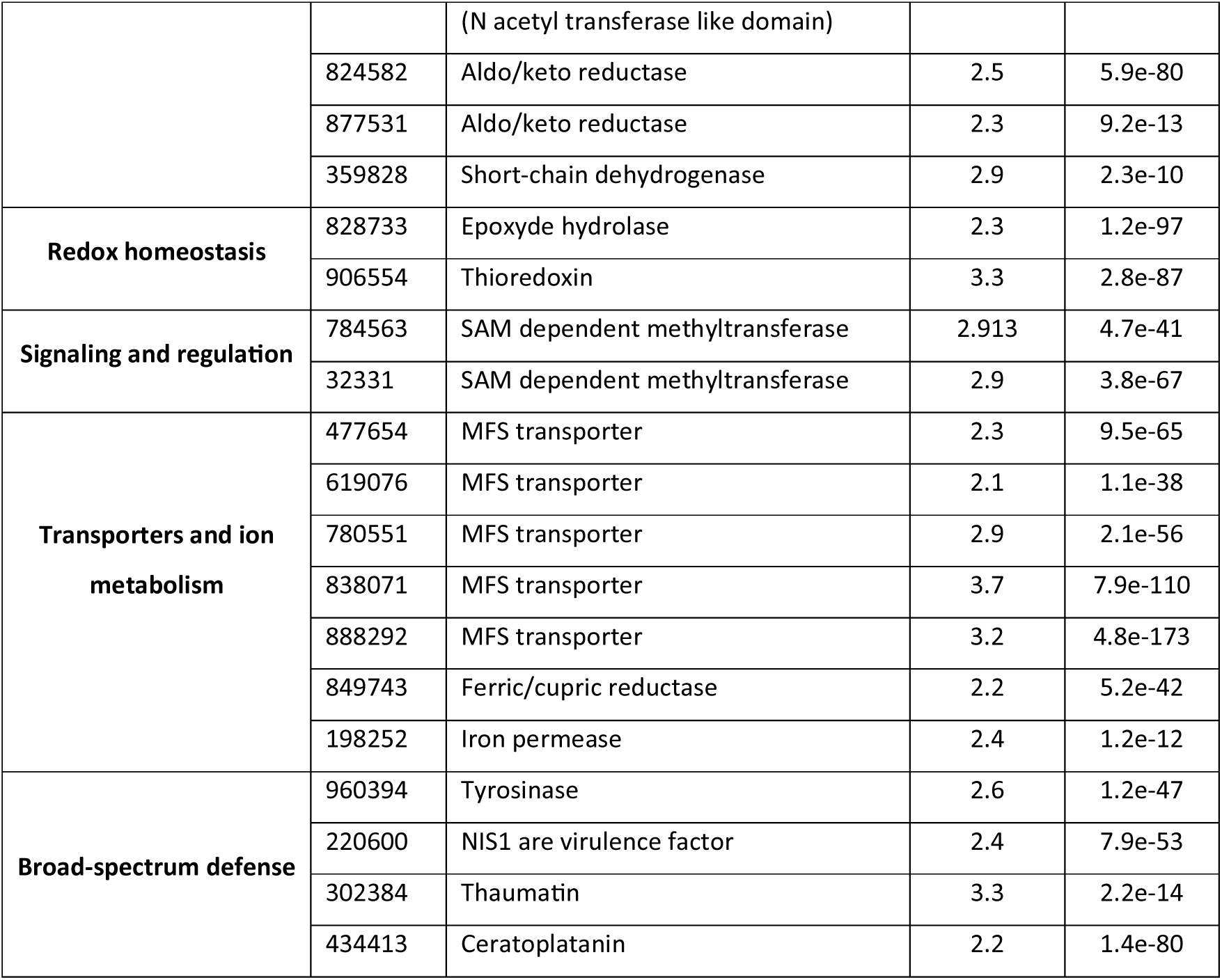
Selection of up-regulated genes in F. betulina CIRM-BRFM 8c0 during the coculture with E. coli. ProtID refer to the accession number of the homologous gene in the genome of F. betulina CIRM-BRFM 1772 on Mycocosm.

### The response to carbon starvation involved interconnected pathways between central and secondary metabolism

We noticed the differential expression of several genes involved in cross-talks between central and secondary metabolism during the coculture. Strikingly, among the up-regulated genes, we identified a Velvet domain (ProtID #496859, induction factor 5), which shared 34.4% identity with the Velvet Complex subunit B of *Laccaria bicolor* S238N-H82 (accession B0CXQ2.1) and *Schizophyllum commune* H4-8 (accession D8PJU0.1). In Fungi, the Velvet Complex is a central regulator of development, metabolism and the stress response, and one of the main central regulators for the synthesis of secondary metabolites (46, 47). For example, it regulates the synthesis of penicillin, lovastatin and the mycotoxin sterigmatocystin (48, 49). The Velvet Complex is a heterotrimeric complex, composed of the subunits VelB, VeA and LaeA, which binds DNA and regulates translation. Few homologs of Velvet Complex genes have been described in Basidiomycota species (*Coprinopsis cinerea, Ganoderma lingzhi,* and *Pleurotus ostreatus*) (47, 50–52). Regarding VelB, it is also related in *Aspergillus nidulans* to sexual reproduction and secondary metabolism (53).

We also found three up-regulated genes involved in the glyoxylate cycle, also known as Tri Carboxylic Acid cycle shunt (ProtID #784186, #290317, and #403655). The aim of the glyoxylate cycle is to produce malate and AcetylCoA as precursors of carbohydrates. The glyoxylate shunt allows fungi to produce carbohydrates from fat in carbon starvation conditions, and AcetylCoA is also used as a precursor by polyketide synthases (54, 55).

We found two up-regulated aldo/keto reductases (ProtID #824582 and #877531) and a short-chain dehydrogenase (ProtID #359828). Aldo/keto-reductases and short-chain dehydrogenases are involved in the breakdown of complex carbon sources (56) and in the cyclization of polyketides during the final steps of the biosynthesis (57).

We identified five up-regulated genes linked to redox homeostasis, among which a predicted epoxide hydrolase (ProtID #822733) and a thioredoxin (ProtID #906554). Lastovetsky et. al. studied the interaction between *Rhizopus macrosporus* (ATCC52813), a saprotrophic fungus (Mucoromycota), and *Mycetohabitans* in endosymbiosis or antagonistic contexts (58). Using transcriptomics, they found an increase in ROS production by the fungus in the antagonistic context, and a decrease in ROS production in the endosymbiosis context (58), highlighting the role of redox homeostasis in the fungal adaptation.

Finally, we found eight up-regulated genes related to signaling and regulation. Among them, two are SAM dependent methyltransferases (ProtID #784563, #32331), also involved in secondary metabolite biosynthesis (59, 60).

### Coculture impact transporters and ion metabolism

Transporters play a key role in fungal nutrition and in the secretion of defense metabolites. We found seven up-regulated transporters, among which five belong to the Major Facilitator Superfamily (MFS) (ProtID #477654, #619076, #780551, #838071, and #888292).

Two genes, a predicted ferric/cupric reductase (ProtID #849743; PF01794), and a predicted iron permease from the FTR1 superfamily (ProtID #198252) were up-regulated and related to iron and cupper uptake. Interestingly, iron is a limiting factor for microbial growth and of particular importance in the outcome of competition between microorganisms (61).

### Defense mechanisms were triggered during the coculture

We identified four broad-spectrum fungal defense mechanisms induced during the coculture. One is related to melanin biosynthesis. The gene #960394 is a tyrosinase (copper monooxygenase), known to be involved in melanin biosynthesis in *Aspergillii* (46). Melanin is reported as a “fungal armor” to protect and survive in harsh conditions through antibacterial activities, thermo-, photo-and radioprotection, as protection against oxidative stress, and as metal chelator (62).

The second, ProtID #220600, is related to necrosis-inducing secreted protein 1 (NIS1). In plant pathogenic fungi, NIS1 are virulence factors that interfere with the basal immunity of the host (63). The ProtID #302384 and ProtID #434413 are related to thaumatins and ceratoplatanins, respectively. Both proteins are secreted by fungi facing adverse growth conditions, such as the absence of easily assimilable carbon (64, 65). Ceratoplatanins are small, hydrophobic proteins that form ordered aggregated layers at hydrophobic/hydrophilic interfaces and contribute to cellulose degradation by disrupting non-covalent bonds in cellulose fibrils (66).

### The coculture triggered the induction of Biosynthetic Gene Clusters

Using ANTISMASH for the identification of Biosynthetic gene clusters, we identified 35 BGCs in the genome of *F. betulina* (Table 2), among which clusters for the biosynthesis of terpenes (17 BGCs) and f-RiPPs-like (9 BGCs) were the most abundant. We identified two hybrid BGCs, one NRPS-T1PKS hybrid, and one fungal-RiPP-Terpene hybrid. No data were available on the predicted metabolites synthesized by these BGCs, except ∂-cadinol that was predicted for three BGCs. Delta-cadinol has been shown to dock on Penicillin-Binding Proteins and is partially responsible of the antibacterial activity of *Cymbopogon martinii* essential oil (67). However, the prediction for the reaction product of the BGCs, and the fungal-RiPP detection should be interpreted with caution (68).

**Table 2:** Numbers of BGC detected with ANTISMASH in the genome of F. betulina. Among them, one BGC is a NRPS-T1PKS hybrid, and one is a fungal-RiPP-Terpene hybrid.

| BGC type | NRPS | Terpene | Fungal-RiPP | T1PKS | Total |
| --- | --- | --- | --- | --- | --- |
| Number of BGC | 6 | 17 | 9 | 5 | 37 |
| Number of genes | 76 | 169 | 121 | 47 | 413 |
| Number of biosynthetic genes | 16 | 39 | 24 | 13 | 92 |

Among the 413 genes identified in the 35 BGCs, we found 15 up-regulated genes that were members of 9 BGCs (Table 3).

**Table 3:**
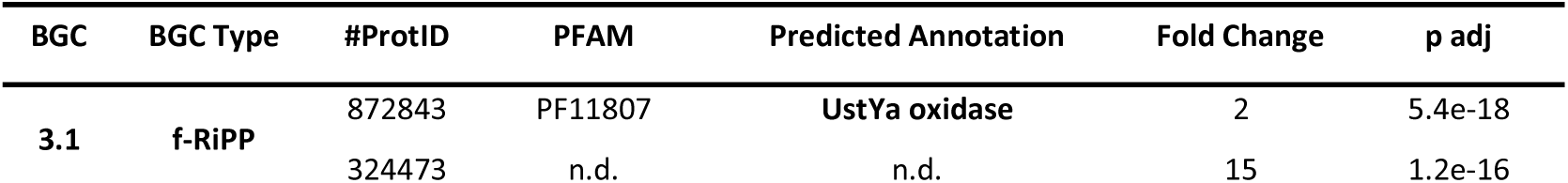
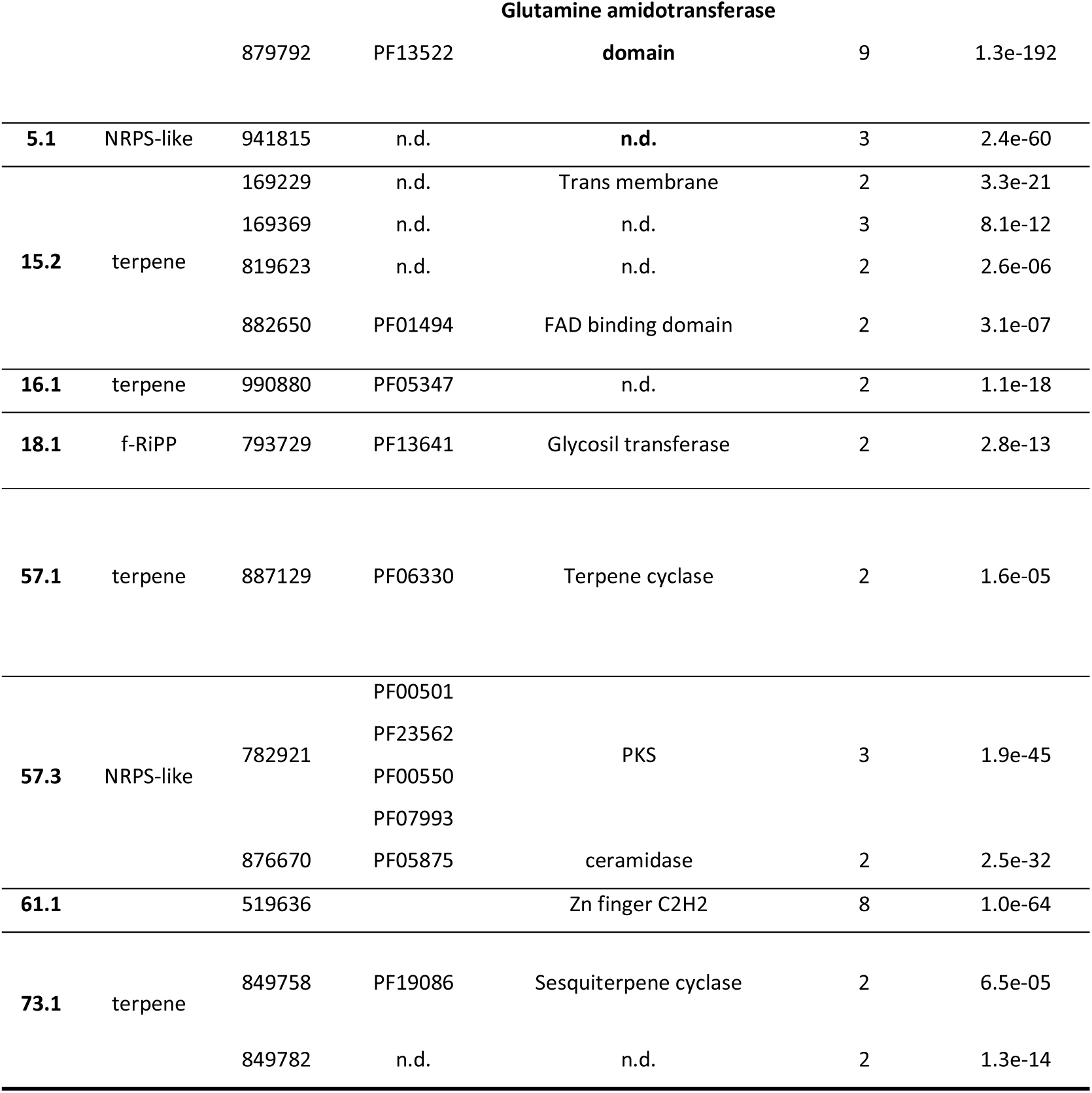
List of the BGC genes up-regulated during the coculture. ProtID refers to the accession number of the homologous gene in the genome of F. betulina CIRM-BRFM 1772 on Mycocosm. (n.d.: no data)

The BGC 3.1, annotated as fungal-RiPP-like, was composed of 18 genes, among which three were up-regulated in our coculture conditions. A BLASTp search against SwissProt showed 27-32% identity between the core synthetic gene (ProtID #872843) and UstYa oxidases involved in the synthesis of fungal RiPPs of the dikaritin family. We found 27% identity with the biosynthetic gene of phomopsin, a mycotoxin from *Diaporthe toxica,* formerly *Diaporthe leptostromiformis,* the agent of lupinosis, 32% identity with the biosynthetic gene of victorin, a mycotoxin from *Bipolaris victoriae, and* 29% identity with biosynthetic genes of asperipin and ustilotoxin B in *Aspergillus niger* NRRL3357. The ustilotoxin and phomopsin have high affinity with tubulin and antimitotic properties (69, 70). Within the same BGC, the protein #879792 contained a Glutamine amidotransferase type 2 domain that could be related to the biosynthesis of β-lactam, a cyclic amid that is found in penicillin (71), and the protein #879792 shared 41.79% identity with a RING finger protein of *Schizosaccharomyces* (accession P87237), which could be involved in expression regulation.

The BGC 61.1, annotated as fungal-RiPP, was composed of 20 genes, among which one was up-regulated in our coculture condition. The core biosynthetic gene of this BGC also shared about 30% (22-35%) identity with UstYa oxidase genes involved in cyclopeptidic mycotoxin biosynthesis in Ascomycota fungi. Besides, the gene #519636 in this BGC contained a Zinc finger domain (C2H2 type) typical of transcription factors.

We also found a terpene cyclase (ProtID #887129), a sesquiterpene cyclase (ProtID #849758) and a polyketide synthase (ProtID #782921) among the up-regulated genes. ProtID #887129 shared respectively 37 and 39% identity with terpene cyclase 29 (accession A0A348B794.1) and 25 (accession A0A348B793.1) of the Polyporales fungus *Postia placenta* Mad-698-R, and 28% identity with an alpha-cuprenene synthase of the phylogenetically related *Agaricales* fungus *Coprinospis cinerea* okayama7#130. ProtID #849758 shared respectively 53 and 54% identity with a muurolene synthase of *C. cinerea* okayama7#130 (accession A8NE23.1) and the sesquiterpene synthase agr3 of the Agaricales *Cyclocybe aegerita* (accession A0A5Q0QU70.1). Cuprenene and muurolene are volatile compounds found in plant essential oils with biological activities. ProtID #782921 shared 29.5% identity with a PKS of the Russulales *Heterobasidion annosum*, and 31% identity with the Non-Reducing PKS tropA of the Ascomycota *Talaromyces stipitatus* ATCC 10500 (accession B8M9J9.1). TropA is responsible for the synthesis of tropolone, non-benzenoid aromatic compounds. Tropolone derivatives have antibacterial and antifungal activities (72, 73).

### Coculture impact globally the fungal metabolic pathways

Finally, our results showed that competitor bacteria strongly impact the fungal metabolic regulation on several aspects, including carbon acquisition, transporters, and redox homeostasis. Thus, in response to the metabolic stress, fungi up-regulates non-specific defense as secondary metabolites biosynthetic genes clusters (Fig. 7). Since a central regulator like the velvet complex is impacted by these competing conditions, the observed impact on several metabolic pathways is rather relevant as consequences. However, some pathways could also be directly impacted by the carbon starvation, and the competition with the bacteria. Thus, regarding carbon acquisition regulation, monomers often are necessary to trigger the secretion of complex CAZymes. The carbon depletion/bacterial competition may not be enough to trigger it, which may avoid the fungi to spend critical resources to synthetize and secrete complex proteins in vain if the substrates are not in the near environment. The up-regulation of transporters could indicate that the fungi “is preparing” to a next to come need of increasing traffic, which is relevant with the up-regulation of nutrients acquisition and metabolites secretions.

It is also interesting to observe that our conditions did not reveal up-regulation on spores’ formation or sexual adaptation that could be regulated by VelB, as showed in other fungi such as in Ascomycota. These results suggest that these pathways are different, or poorly annotated, in Basidiomycota.

It is also interesting to observe that despite the up-regulation of VelB, numerous unknown genes are down-regulated. Could this be a consequence? Or should we look for different pathways directly regulated, and negatively impacted during the carbon competition? Thus, no specific interactions mechanisms, as it could have been expected have emerged. This could mean that the response of fungi to bacterial competitor might be closely strains and context specific, and complex. It would be very interesting to compare the fungal regulations in response to different bacterial environmental strains and specific stresses, like direct oxidative stress, or glucose depletion to decipher which pathways, if any, are specific or not, of such conditions.

## Conclusion

This study demonstrates that bacterial competition can act as a powerful trigger of metabolic adaptation in the brown-rot fungus *F. betulina* CIRM-BRFM860. Co-culture with *E. coli* ATCC25922 rapidly depleted the available carbon resources, resulting in carbon starvation for the fungus and a marked reprogramming of fungal metabolism. The fungal response involved the induction of genes associated with carbon acquisition, nutrient transport, redox homeostasis, broad-spectrum defense mechanisms, and specialized metabolism. Notably, the up-regulation of a VelB homolog highlights the potential involvement of the Velvet regulatory complex in coordinating the fungal response to bacterial competition and metabolic stress.

Our results suggest that the production of secondary metabolites in *F. betulina* is driven, at least in part, by resource competition. More broadly, they indicate that the ecological pressures encountered during microbial competition can stimulate the expression of otherwise poorly expressed biosynthetic pathways. The induction of genes belonging to multiple biosynthetic gene clusters, including terpene, polyketide, and fungal RiPP pathways, further supports the potential of co-culture approaches for accessing underexplored fungal metabolic diversity.

Taken together, these findings improve our understanding of fungal-bacterial interactions in wood-decaying ecosystems and establish nutrient competition as a key driver of metabolic regulation in basidiomycetes. They also highlight co-culture-based strategies as promising tools for activating silent biosynthetic pathways and accelerating the discovery of novel fungal natural products with potential biotechnological and pharmaceutical applications.

## Acknowledgement

All authors warmly thank the technical staff for their routine lab work and the administrative staff for their support. The authors acknowledge Charlotte SIMMLER, co-responsible for the Metabolomics and Natural Product chemistry (MSN) facility at IMBE, for the NMR acquisitions of biomarker#6 and its data sharing. The authors acknowledge Plateforme GeT-Biopuces, TBI, Université de Toulouse, CNRS, INRAE, INSA, Toulouse, France and particularly Delphine Labourdette for their work on the transcriptomic raw data. The authors acknowledge Lise Molinelli for the help regarding the synthesis of piptamine.

## Author’s contribution

Fundings acquisition: QA, MNR

Experimental design: QA, IG, MNR

Chemical analyses and Metabolomics: QA, DN, SG, PV

Chemical synthesis: MR, AD

Antibacterial activity: QA, MM

RNA extraction and Transcriptomic data analysis: QA, JL, ED, MNR

Manuscript draft: QA, MNR

Manuscript editing: QA, ML, SG, MNR

## Fundings

The project leading to this publication has received fundings from the Excellence Initiative of Aix-Marseille University – A*MIDEX, a French “Investissements d’Avenir” programme and is part of the Institute of Microbiology, Bioenergies and Biotechnologies – IM2B (AMX-19-IET-006)

This research has been financially supported by INRAE and CARNOT 3BCAR.

## Competing interests

The authors declare that they have no competing interests.

